# Discovery and Characterization of Interleukin-4-Specific Affibodies for Affinity-Controlled Protein Release and Macrophage Polarization

**DOI:** 10.64898/2026.05.07.723637

**Authors:** Jonathan Dorogin, Astha Lamichhane, Andy J. Huang, Justin E. Svendsen, Morrhyssey A. Benz, Shreya A. Raghavan, Marian H. Hettiaratchi

**Affiliations:** Department of Bioengineering, Phil and Penny Knight Campus for Accelerating Scientific Impact, University of Oregon; Department of Biomedical Engineering, Texas A&M University; Department of Chemistry and Biochemistry, University of Oregon; Institute of Molecular Biology, University of Oregon

**Keywords:** Interleukin-4, affibody, hydrogel, protein delivery, affinity interactions

## Abstract

Interleukin-4 (IL-4) is a key immunoregulatory cytokine that promotes type 2 inflammation, drives macrophage polarization toward an anti-inflammatory M2 phenotype, and supports tissue repair. However, clinical translation of IL-4 therapies to modulate the immune response is limited by the need for precise control over its delivery to avoid immune dysregulation. Here, we report an affinity-based strategy to modulate IL-4 delivery and bioactivity using engineered affibody proteins. A yeast surface display library was screened via magnetic- and fluorescence-activated cell sorting to identify two IL-4-specific affibodies with moderate binding affinities (dissociation constants, K_D_ = 459 and 141 nM). Circular dichroism confirmed expected alpha-helical folding, and biolayer interferometry characterized the kinetics of IL-4 binding. Structural modeling using AlphaFold3 and RosettaDock and molecular dynamics simulations using GROMACS predicted distinct binding sites for each IL-4-specific affibody on the IL-4 protein and suggested potential interference with receptor complex formation. Bioactivity studies using murine bone marrow-derived macrophages demonstrated that IL-4 complexed with affibodies maintained *Ym1* gene expression but significantly reduced Ym1 protein levels, indicating partial inhibition of IL-4 signaling. To enable controlled cytokine delivery via affinity interactions, affibodies were conjugated to polyethylene glycol maleimide (PEG-mal) hydrogels, which were loaded with IL-4. Affibody-conjugated hydrogels achieved high IL-4 loading efficiency (>90%) and exhibited sustained release over 7 days. Increasing affibody-to-IL-4 ratios significantly reduced both the rate and total amount of cytokine release. Overall, this work establishes IL-4-specific affibodies as versatile tools for tuning cytokine presentation and modulating bioactivity and provides a promising approach for regulating inflammatory responses and advancing cytokine-based therapies with improved temporal control.

**Graphical Abstract:** 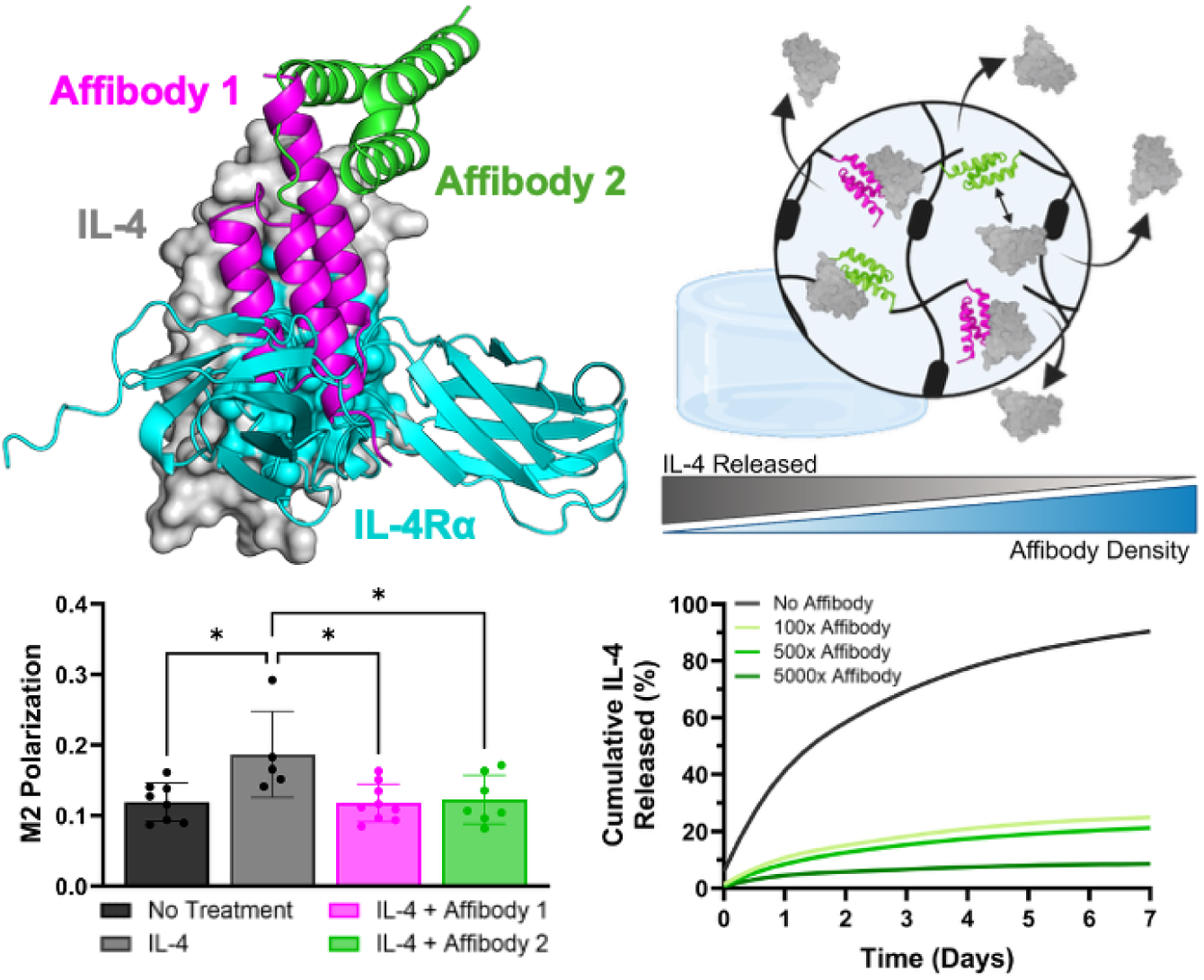

## Introduction

The inflammatory response is a fundamental defense mechanism that protects the body from infection, injury, and other harmful stimuli.^1^ It involves the highly coordinated interplay of numerous immune cells and signaling molecules, which participate in both stimulatory and regulatory functions to fight infection and repair damaged tissues without causing excessive inflammation. Interleukin-4 (IL-4) is a particularly dynamic cytokine that plays a critical role in coordinating the immune response to resolve inflammation, characterized by the differentiation of naïve T cells into T helper 2 (Th2) cells, polarization of macrophages into the M2-like spectrum towards an anti-inflammatory phenotype, suppression of pro-inflammatory factor production, and stimulation of anti-inflammatory factor production.^2,3^ These actions are regulated through the interaction of IL-4 with the IL-4 receptor α (IL-4Rα) and to a lesser extent IL-13 receptor α (IL-13Rα), which ultimately activates the transcription of signal transducer and activator transcript 6 (STAT6).^4^ In myeloid cells like macrophages, IL-4 signaling triggers the receptor expression for mannose and chitinase-like proteins like Ym1, a hallmark of pro-resolving macrophage polarization primed for wound healing and tissue remodeling. IL-4 signaling also impacts other cell populations, including stimulating the expression of vascular cell adhesion molecule-1 (VCAM-1) on endothelial cells to promote immune cell homing to sites of inflammation, promoting allergic airway responses through increased smooth muscle cell contractility, and providing neuroprotection after injury to the brain.^5–7^ The pro-regenerative effects of IL-4 during tissue repair, which include cytokine and chemokine secretion, angiogenesis, phagocytosis, and extracellular matrix remodeling, are largely mediated by the T helper 2 pathway and executed by M2 macrophages.^8^

Given the role of IL-4 in mitigating inflammation, reestablishing immune homeostasis, and supporting tissue repair, exogenous IL-4 delivery has been investigated as a means to stimulate M2-like macrophage polarization,^9,10^ improve wound healing,^11,12^ attenuate cognitive deficits after traumatic brain injuries,^13,14^ and reduce bone resorption.^15,16^ However, the short half-life and relative instability of this cytokine necessitates repeated daily injections or topical administration to elicit long-term effects.^11 12^ Alternatively, diffusion-mediated IL-4 release from nanoparticles, microparticles, and bulk hydrogels,^7,17,18^ local presentation of biotinylated IL-4 using biotin-streptavidin affinity interactions,^10,19^ and the delivery of mesenchymal stromal cells genetically engineered to overexpress IL-4^15^ can provide sustained IL-4 delivery to stimulate macrophages and other cell populations. In the native immune response, the temporal dynamics of IL-4 signaling are tightly regulated, ensuring that an appropriate balance of pro- and anti-inflammatory signaling molecules can be maintained during both homeostatic conditions and an active immune response to injury or infection. Prolonged exposure to IL-4 can lead to immune dysregulation, allergic inflammation, and asthma.^20–22^ Thus, simply extending IL-4 delivery over a long period of time without control over the rate and amount of cytokine released may not be sufficient to stimulate the necessary transition from pro-inflammatory to anti-inflammatory responses.

To provide better tunability over protein delivery, we and others have developed affinity-based delivery vehicles that control protein release and bioactivity through natural or engineered non-covalent affinity interactions between the target protein and biomaterial.^23–29^ Several extracellular matrix molecule-derived polymers such as heparin and fibronectin display intrinsic affinities towards heparin-binding proteins, including bone morphogenetic protein-2,^30^ vascular endothelial growth factor,^31^ and platelet-derived growth factor.^32^ These polymers can be modified to adjust their degree of sulfation or other charged functional groups to manipulate their affinity interactions and tune the local availability of delivered proteins.^31,33,34^ Affinity interactions with aptamers,^35^ peptides,^36^ antibody fragments,^37^ and antibody mimetics such as affibodies and nanobodies^26^ that are engineered to bind to target proteins with high specificity and variable affinities can provide increased tunability over protein release rates. Both natural and engineered affinity interactions have also been shown to affect interactions between target proteins and their receptors in a dose-dependent manner, functioning as vehicles for modulating protein bioactivity with spatiotemporal control.^26,30^

Here, we demonstrate an affinity-based strategy to control the delivery and bioactivity of IL-4 with the goal of providing a tunable platform to finely manipulate IL-4 presentation and its effects on macrophages. We screened a yeast surface display library of affibody proteins to identify two moderate-affinity affibodies that specifically bind to IL-4. We leveraged the computational protein design tools AlphaFold3 and RosettaDock, as well as GROMACS molecular dynamics software, to predict the mechanisms of binding between the affibodies, IL-4, and IL-4Rα. We investigated the effect of IL-4-specific affibodies on IL-4-mediated M2 polarization of murine bone marrow-derived macrophages, demonstrating the potential of these affibodies to modulate IL-4 activity and validating the computationally predicted interactions between the IL-4-specific affibodies and IL-4. Finally, we conjugated IL-4-specific affibodies to polyethylene glycol maleimide (PEG-mal) to impart engineered affinity to PEG-mal hydrogels, achieving IL-4 release profiles that could be tuned by changing the ratio of affibodies to IL-4. Taken together, this work establishes a new platform for tuning IL-4 delivery and activity for the modulation of type 2 inflammation.

## Materials and Methods

### Yeast cell culture

Yeast cell culture was performed as previously described.^25^ Briefly, an affibody-expressing *S. cerevisiae* EBY100 yeast surface display library containing 4 × 10^8^ unique affibody sequences was cultured in selective growth media (16.8 g sodium citrate dihydrate, 3.9 g citric acid, 20.0 g dextrose, 6.7 g yeast nitrogen base, 5.0 g casamino acids, 1 mg ciprofloxacin and 100 mg ampicillin in 1 L reverse osmosis (RO) water) with orbital shaking at 30 °C for 20 hours to a concentration of 10^8^ cells/mL. At least 10× library diversity was transferred to a new vessel, and surface expression of affibodies was induced using selective induction media (10.2 g sodium phosphate dibasic heptahydrate, 8.6 g sodium phosphate monobasic monohydrate, 19.0 g galactose, 1.0 g dextrose, 6.7 g yeast nitrogen base, 5.0 g casamino acids, 1 mg ciprofloxacin and 100 mg ampicillin in 1 L RO water) with orbital shaking at 30 °C for 20 hours.

### IL-4-specific affibody identification using magnetic- and fluorescence-activated cell sorting

The yeast surface display library was enriched for IL-4-specific affibodies using magnetic-(MACS) and fluorescence-activated cell sorting (FACS) by adapting a previously described protocol.^25^ For MACS, carboxylic acid surface-modified magnetic beads (COOH beads) (Invitrogen) were conjugated with 0.1% (w/v) bovine serum albumin (BSA) in phosphate-buffered saline (PBS) or 0.05 M tris for negative sorts or 33 pmol of IL-4 (Peprotech, Inc.) for the positive sort using carbodiimide chemistry. Affibody-expressing yeast were washed in 0.1% (w/v) BSA in PBS to remove the induction media and 10× library diversity was incubated with BSA-conjugated COOH beads for 2 hours at 4 °C with rotation. Beads were exposed to a magnet, and the unbound yeast solution was gently removed and transferred to a tube containing tris-conjugated magnetic beads for 2 hours of incubation at 4 °C with rotation. The magnet exposure was repeated, and the unbound yeast solution was transferred to a tube containing IL-4-conjugated magnetic beads for 2 hours at 4 °C with rotation. The IL-4-positive magnetic beads were then resuspended in 0.1% (w/v) BSA in PBS. The BSA, tris, and IL-4 beads were diluted 100 and 2000 times, and 10 µL of each solution was plated on selective growth plates (16.8 g sodium citrate dihydrate, 3.9 g citric acid, 16 g bacto agar, 20 g dextrose, 6.7 g yeast nitrogen base, 5 g casamino acids, RO water) and incubated for 36 hours at 30 °C. The number of yeast removed by the BSA and tris bead negative selections was estimated by counting the colonies on the BSA and tris bead selective growth plates. The new yeast library diversity after each round of MACS was estimated by counting the number of colonies on the IL-4 bead plates, while the ratio of positive-to-negative binders was calculated by dividing the number of colonies on the IL-4 bead plates by the sum of colonies on the negative BSA and tris bead plates. The new library diversity was used to determine the quantity of yeast needed for subsequent sorts.

Following 5 rounds of MACS, the enriched yeast display library was subjected to FACS.^25^ 40 × 10^6^ affibody-expressing yeast cells were resuspended in 50 µL of 0.1% (w/v) BSA in PBS containing 1 µM biotinylated IL-4 (Acro Biosciences) and 1.25 µL of anti-c-Myc mouse monoclonal antibody (9E10; BioLegend) and rotated at 4 °C for 90 minutes. The cells were washed twice with 500 µL of 0.1% (w/v) BSA in PBS and then resuspended in 50 µL of a secondary antibody solution containing 17.3 nM goat anti-mouse IgG AlexaFluor™ 647 (Invitrogen) and 540 nM of AlexaFluor™ 488 streptavidin conjugate (Invitrogen) in 0.1% (w/v) BSA in PBS for 30 minutes at 4 °C in the dark. The cells were washed twice with 500 µL of 0.1% (w/v) BSA in PBS and then resuspended in 1000 µL of 0.1% (w/v) BSA in PBS for FACS using a SH800 Cell Sorter (Sony Biotechnology). Controls to set FACS gates included yeast cells without any labeling, yeast cells labeled with secondary solution only, and yeast cells labeled with anti-c-Myc antibody and secondary solution. During sorting, yeast cells were collected from the double positive (488+/647+) quadrant, resuspended in selective growth media, and expanded at 30 °C to approximately 5 × 10^6^ cells/mL before inoculating selective growth plates at 30 °C for 36 hours to isolate single colonies.

Single colonies were expanded in selective growth media to a cell density of approximately 10^7^ cells/mL. The yeast plasmids were isolated using an Easy Yeast Plasmid Isolation Kit (Clontech), the affibody sequences were amplified using PCR with custom primers for the affibody inserts (forward primer (5′-CCCTCAACAACTAGCAAAGG-3′) and reverse primer (3′-ATGTGTAAAGTTGGTAACGGAACG-5′)), and the products were purified using a DNA Clean and Concentrator Kit (ZymoGen). Sanger sequencing was performed by Genewiz (Azenta Life Sciences).

Flow cytometry was used to confirm binding of single colonies of affibody-expressing yeast to IL-4. The FACS protocol was modified to use 1 × 10^6^ yeast cells on an Accuri C6 Plus flow cytometer (Becton Dickinson). Binding to IL-4 was quantified by comparing the number of double positive cells (AF647+/AF488+) in samples incubated with anti-c-Myc antibody, biotinylated IL-4, and secondary antibody solution to samples incubated with only the anti-c-Myc antibody and secondary antibody solution. Two unique IL-4-specific affibodies (Affibody 1 and Affibody 2) were identified.

### Solid-phase peptide synthesis of affibodies

IL-4-specific affibody sequences were modified to include a penultimate cysteine and a C-terminal glycine and synthesized using solid-phase peptide synthesis (SPPS) on Fmoc-Gly-Wang resin (CEM Corporation) with a 0.602 mmol/g loading capacity using a Liberty Blue 2.0 microwave peptide synthesizer (CEM Corporation), as previously described.^38^ 0.2 M amino acid (CEM Corporation), 10% (v/v) pyrrolidine (Sigma Aldrich) deprotection, 1 M N,N’-diisopropylcarbodiimide (DIC) (Oakwood Chemical), and 1 M Oxyma (CEM Corporation) solutions were prepared in dimethylformamide (DMF). After synthesis, resin-bound affibodies were washed with dichloromethane (DCM) twice and resuspended in a solution containing a 1:1:1:1:36 ratio of ddH_2_O, triisopropyl silane (Oakwood Chemical), 2,2’-(ethylenedioxy)diethanethiol (Tokyo Chemical Industry), thioanisole (Oakwood Chemical), and trifluoroacetic acid (TFA) (Oakwood Chemical) for 40 minutes at 42 °C for cleavage from the resin.^39^ The solution was vacuum-filtered, and the filtrate was purified via precipitation and three rounds of centrifugation (1200 rcf for 5 minutes) using cold diethyl ether. The resulting slurry was dried overnight in a vacuum desiccator and transferred to −20 °C for storage.

To purify the synthesized affibodies, the crude product pellets (∼650 mg) were dissolved in 3% (v/v) acetonitrile and 0.1% (v/v) TFA in ddH_2_O and subjected to high-performance liquid chromatography (HPLC) (CEM) using 19 x 150 mm C18 Waters column by running a 3-75% (v/v) acetonitrile gradient in ddH_2_O with 0.1% (v/v) TFA. Affibody_1 was eluted from the column at approximately 61% (v/v) acetonitrile in ddH_2_O, while Affibody_2 was eluted from the column at 63.5% (v/v) acetonitrile in ddH_2_O. Approximately 350 mg of each purified affibody was collected in a 50 mL conical tube, frozen at −80 °C, lyophilized at −104 °C and 50 mTorr for 24 hours using a benchtop lyophilizer (SP Scientific). Purified affibodies were stored at −20 °C until use.

### Soluble affibody characterization

The size, secondary structure, and IL-4-binding interactions of the IL-4-specific affibodies were characterized using matrix-assisted laser desorption/ionization time of flight (MALDI-TOF) mass spectrometry, circular dichroism, and biolayer interferometry (BLI), respectively. For MALDI-TOF, 1 µL of 1 mg/mL affibody in 3% (v/v) acetonitrile and 0.1% (v/v) TFA in ddH_2_O and 1 µL of matrix solution (10 mg/mL α-cyano-4-hydroxycinnamic acid (Sigma Aldrich) in 50% (v/v) acetonitrile and 0.1% (v/v) TFA in ddH_2_O) were deposited onto a stainless steel target plate (Bruker). The products were analyzed on a MALDI-TOF Smart LS system (Bruker) using 200 event counts and fitted to a 3-20 kDa protein standard (Bruker). Baseline noise was background subtracted in the Bruker software.

A Jasco J-815 circular dichroism spectropolarimeter was used to measure affibody secondary structure. Affibodies were reconstituted in 10 mM tris buffer, pH 8 at approximately 0.5 mg/mL. 200 µL of affibody solution was loaded into a quartz cuvette (Starna Cells, Inc.) with a 1 mm path length. Circular dichroism spectra were obtained between 190-250 nm and normalized to mean residue molar ellipticity.^40–42^

Binding between IL-4 and IL-4-specific affibodies was measured using a GatorPlus biolayer interferometer (Gator Bio). Streptavidin-coated (SAXT) probes were loaded with 1 nM of biotinylated IL-4 (Acro Biosystems) in 0.05% (v/v) Tween 80 and 0.2% (w/v) BSA in PBS (buffer solution), quenched with 20 µg/mL of biocytin (Caymen Chemicals) in buffer solution, and then allowed to associate to and dissociate from 15.6-4000 nM of IL-4-specific affibodies in buffer solution for 120 seconds per step. The dissociation constants (K_D_), on-rate constants (*k_on_*), and off-rate constants (*k_off_*) were calculated for each IL-4-affibody binding interaction by curve fitting the wavelength shifts using a 1:1 binding fit model in the GatorOne software (Gator Bio).

### Computational predictions of IL-4-affibody binding

IL-4-specific affibodies were folded and docked to IL-4 *in silico* using AlphaFold3 multimer prediction.^43^ Sequences for IL-4, Affibody 1, and Affibody 2 were input into AlphaFold3 and recycled three times to generate the five best binding predictions. Complexes of affibodies bound to IL-4 were energy minimized using Rosetta FastRelax scripts to create bound conformations with the lowest energies for interface screening. The highest ranked structural prediction for each affibody-IL-4 complex was aligned onto the x-ray crystallography structure of IL-4 binding to its receptor, IL-4 receptor-α (IL-4Rα, PDB: 1IAR) to enable direct comparisons of predicted bound interfaces. Energies of bound complexes were measured using Rosetta FastRelax script metrics to compare predicted interface stabilities of energy minimized bound states.^44^ 5000 rounds of docking were performed using the RosettaDock protocol to model IL-4 binding to IL-4Rα, Affibody 1, and Affibody 2.^45,46^ The binding interfaces were simulated across the entire surface of IL-4, starting from the IL-4 receptor-α binding epitope on IL-4. Interface energy scores as a function of root mean square deviation (RMSD) from the initial binding interface were plotted to determine the interface stabilities and selectivity for each IL-4 affibody.^47^

Molecular dynamics simulations were performed on the Rosetta-relaxed structures of IL-4 bound to Affibody 1 or Affibody 2 using GROMACS 2023.4. All simulations were performed using the CHARM36m force field with TIP3P waters with steepest energy descent gradients to create energy minimized systems. We ran four independent 100 ns trajectories for each IL-4-affibody binding interaction. MDAnalysis software packages were used to analyze the results, calculating the all α-carbon RMSD, change in solvent accessible surface area (ΔSASA) of the IL-4-affibody binding interaction, ΔSASA of the unbound vs. bound hydrophobic surface area in the system, ΔSASA of the unbound vs. bound hydrophilic surface area in the system, and affibody residue root mean square fluctuation (RMSF). All scripts used to set up and run the molecular dynamics simulations are available at https://github.com/harmslab/setup_md.

### Macrophage cell culture and exposure to affibodies

Murine immortalized bone marrow-derived macrophages (iBMDMs) were maintained in complete Dulbecco’s Modified Eagle Medium (DMEM) supplemented with 10% heat inactivated fetal bovine serum (FBS), and 1% (v/v) antibiotics and antimycotics. Cells were routinely cultured and sub-cultured at 37 °C with 5% (v/v) carbon dioxide. To assess how IL-4-specific affibodies affected macrophage polarization, iBMDMs were seeded at sub-confluent densities of 3500 cells/mL in 6-well plates and allowed to adhere overnight. Prior to any treatment, microscopic confirmation of live cells was obtained. Affibodies were either added to cells on their own or complexed with IL-4 protein at a ratio of 1:5000 cytokine to affibody for 30 minutes prior to cell exposure, similar to previous experiments.^25^ The experimental groups were as follows: no treatment, IL-4 (50 ng/mL), Affibody 1 (250 μg/mL), Affibody 2 (250 μg/mL), IL-4 complexed with Affibody 1 (1:5000), and IL-4 complexed with Affibody 2 (1:5000). Cells were maintained in these culture conditions for 3 subsequent days following treatment with live cell phase contrast microscopy tracking cell growth.

### Gene expression analysis

After 3 days of treatment, iBMDMs were lysed with Buffer RLT, and RNA was extracted following manufacturer’s protocols using a RNeasy extraction kit (Qiagen, Hilden, Germany). RNA quantity and purity were assessed using a Nanodrop 2000 spectrophotometer, ensuring ratios of 260/280 and 260/230 absorbances greater than 2. cDNA was synthesized using a high-fidelity cDNA transcription kit using a QuantStudio (Applied Biosystems, Foster City, CA). Following reverse transcription, RT-qPCR was performed using SYBR as a fluorescent readout on the QuantStudio qPCR system (Thermo Fisher Scientific), following well established protocols.^48^ Gene expression for *Ym1 (Chi3l3)* and the housekeeping control *Gapdh* was measured (primer sets are provided in **Table 1**). Changes in gene expression were quantified using the 2^−ΔΔCT^ method, with *Gapdh* as the housekeeping gene. qPCR experiments were run in triplicate, with three biologically independent samples.

**Table 1.**
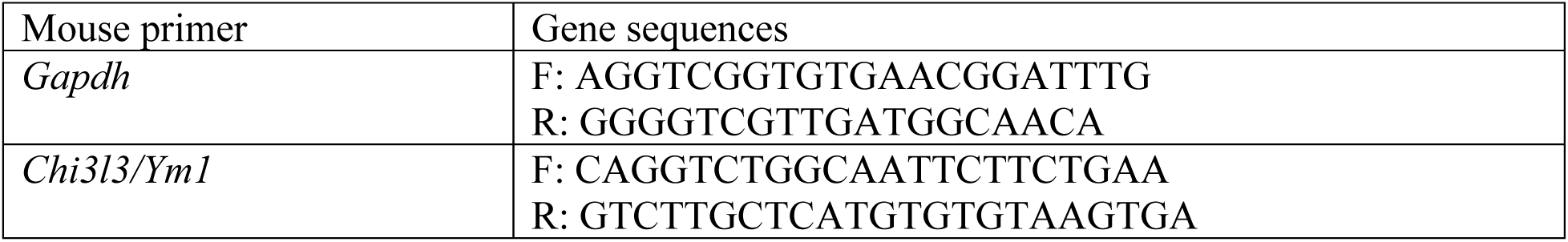
Primers used in RT-qPCR analysis of gene expression.

### Protein expression analysis

To evaluate Ym1 protein expression in iBMDMs, cells were fixed after 3 days of treatment in 4% (v/v) neutral buffered formaldehyde and then quenched and washed with 0.1 M glycine. Cells were permeabilized and blocked with 5% (v/v) goat serum supplemented with 0.5% (v/v) Triton-X. Cells were then incubated with PE-fluorophore-conjugated Ym1 antibody at a 1:100 dilution (Abcam Cat #ab211621) overnight at 4°C. Cells were washed and mounted in Prolong Gold anti-fade reagent. Ym1 fluorescence was imaged with a Nikon AX-R confocal inverted fluorescence microscope (Nikon Instruments Inc., Melville, NY) with gain and amplification settings maintained constant across all experimental conditions.

Quantitative analysis was performed using the corrected total fluorescence (CTF) method in FIJI according to previously established protocols.^49^ For each image, integrated density, area, and mean background fluorescence were measured for each region of interest (ROI). CTF was calculated using the following formula: CTF = Integrated Density – (Area of ROI × Mean Background Fluorescence). To account for variations in cell numbers, CTF was divided by the number of nuclei (DAPI-positive cells) in each image, yielding an average CTF per cell. Each experimental condition was replicated across multiple biological samples (n ≥ 3).

### Hydrogel synthesis and IL-4 release

100 µL 5% w/v affibody-conjugated polyethylene glycol maleimide (PEG-mal) hydrogels were synthesized in 2 mL microcentrifuge tubes under sterile conditions as previously described.^50,51^ All reagents were sterile filtered using 0.22 µm polyethersulfone (PES) syringe filters and handled in a biosafety cabinet. Briefly, 5 mg of 20 kDa, 4-arm PEG-maleimide (Laysan Bio) and 0.66, 3.3., or 33.1 nmol of either Affibody 1 or Affibody 2 were separately dissolved in PBS pH 6.9 and then mixed for 30 minutes at room temperature to react the cysteines on the affibodies with the maleimides on the PEG-mal. The remaining maleimide functional groups on the PEG-mal were then crosslinked using 492 nmol of dithiothreitol (DTT) (Gold Biotechnology) in PBS pH 6.9 at room temperature for an additional 30 minutes. The hydrogels were washed with 1 mL of fresh PBS exchanged 3 times every 12 hours to remove unreacted DTT and affibodies.

Loading and release of IL-4 were also performed as previously described.^38,50^ PEG-mal hydrogels were loaded with 20 µL of 5 µg/mL of IL-4 (100 ng total, equating to 100, 500, and 5000 times molar excess of affibodies to IL-4) overnight with rotation at 4 °C to allow the IL-4 to diffuse into the hydrogel. 200 µL of 0.1% w/v BSA in PBS were added to the hydrogels and immediately removed to determine the amount of IL-4 loaded in the hydrogels. 900 µL of 10% v/v fetal bovine serum (FBS) in PBS were added to the hydrogels to initiate IL-4 release, and 200 µL aliquots were collected and replaced with 200 µL of fresh 10% (v/v) FBS in PBS at 4 hours and 1, 2, 3, 5, and 7 days. IL-4 concentrations were measured using an IL-4-specific enzyme linked immunosorbent assay (ELISA; R&D Systems).

## Results

### Yeast surface display identified two unique IL-4-specific affibodies

Five rounds of MACS were required to sufficiently enrich the yeast surface display library for IL-4 binders. The diversity of the yeast display library progressively decreased over subsequent rounds of MACS, resulting in a final diversity of approximately 4.3×10^5^ unique affibody-displaying yeast with a ratio of 0.413 positive-to-negative IL-4 binders (**Figure 1A**). Following MACS, two rounds of FACS were performed to further enrich the library for IL-4-specific affibodies, and at least 10,000 cells were collected from the AF647+/AF488+ double positive quadrant in each round. Isolating monoclonal yeast and evaluating their binding to IL-4 using flow cytometry identified two unique affibodies that demonstrated binding to IL-4, shown by the increase in AF647+/AF488+ double positive cells with anti-C-myc and biotinylated IL-4 labeling compared to anti-C-myc labeling alone: Affibody 1 (**Figure 1B**) and Affibody 2 (**Figure 1C**). Compared to previously published affibodies discovered from this yeast surface display library,^26,50,52^ this discovery process required more screening steps. It is possible that the relative instability of IL-4, which has a half-life of 19 mins *in vivo*,^53^ reduced the efficacy of the MACS and FACS screening protocols.

**Figure 1.**
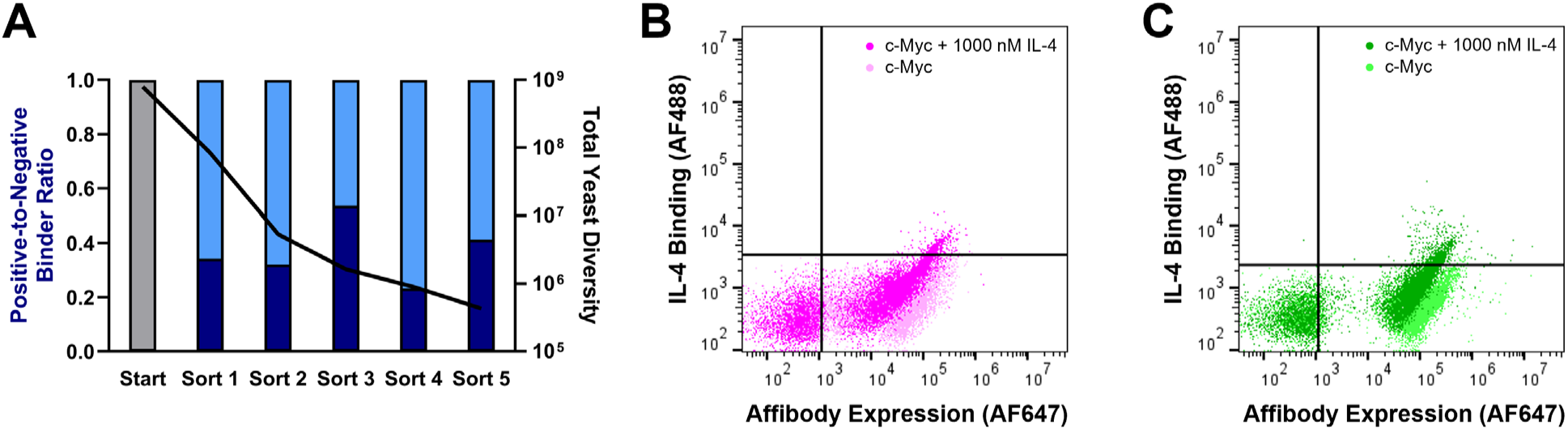
Identification of IL-4-specific affibodies using MACS and FACS. A) Changes in positive-to-negative IL-4 binder ratio (left axis, bar plot) and total yeast library diversity (right axis, black line) over five MACS sorts. Flow cytometry plots of B) Affibody 1 and C) Affibody 2 depicting affibody expression on the x-axis (AF647+) and affibody binding to 1000 nM of IL-4 on the y-axis (AF488+). Yeast displaying affibody binding to IL-4 (c-Myc + 1000 nM IL-4) was assessed in comparison to yeast incubated with anti-c-Myc antibody without IL-4 (c-Myc).

Mass spectrometry revealed a single peak for Affibody 1 at 6749 Da and a single peak for Affibody 2 at 6595 Da (**Figure 2A**), which was comparable to their expected masses of 6748 Da and 6597 Da, respectively. Circular dichroism confirmed that the secondary structures of Affibody 1 and Affibody 2 were α-helical with characteristic troughs at 208 and 222 nm and a peak at 193 nm (**Figure 2B**).^40^ BLI measurements of affibodies binding to IL-4 yielded an equilibrium dissociation (K_D_) constant of 459 nM for Affibody 1 binding to IL-4 (**Figure 2C**) and 141 nM for Affibody 2 binding to IL-4 (**Figure 2D**) with similar on and off rate constants (**Table 2**). In comparison, the interaction between IL-4 and its receptor IL-4Rα has an equilibrium dissociation constant of approximately 1 nM,^54,55^ which is several orders of magnitude stronger than the binding interactions of either of the IL-4-specific affibodies to IL-4, suggesting that these affibodies are both moderate-affinity binders for IL-4. It is important to note that IL-4 signaling proceeds when IL-4 forms a complex with IL-4Rα and either the γ-complex of another IL-4Rα receptor or with a complete IL-13Rα receptor and that these secondary binding interactions have weaker affinities with equilibrium dissociation constants of 559 nM and 487 nM, respectively.^55^

**Figure 2.**
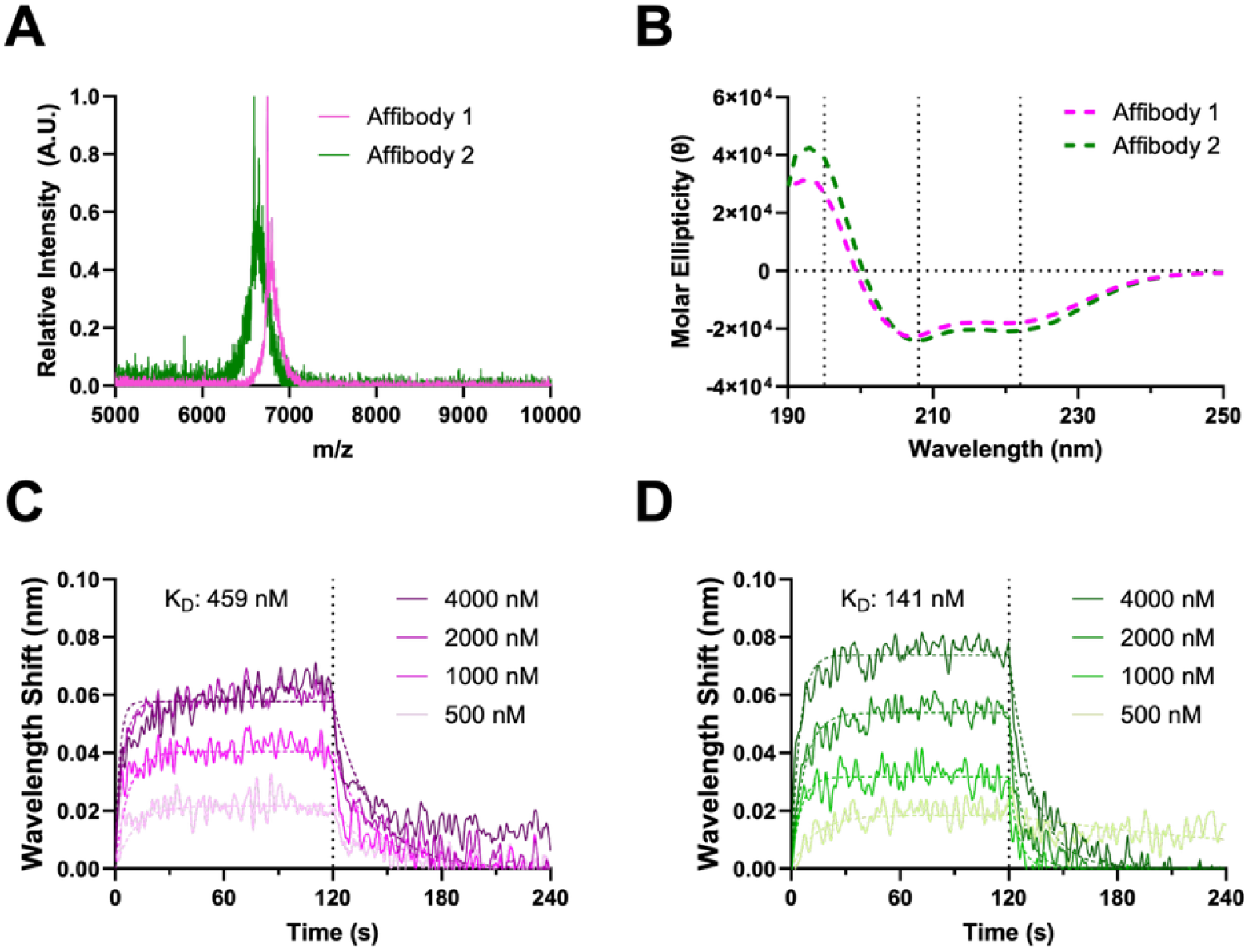
Biochemical characterization of IL-4-specific affibodies. A) MALDI-TOF mass spectrometry spectra of IL-4-specific affibodies, depicting a single peak at 6749 Da for Affibody 1 and a single peak at 6595 Da for Affibody 2. B) Circular dichroism spectra for Affibody 1 and Affibody 2, indicating α-helical secondary structure. BLI was used to calculate equilibrium dissociation constants for Affibody 1 and Affibody 2 binding to IL-4. C) BLI association and dissociation curves for 500-4000 nM of Affibody 1 binding to 1 nM of biotinylated IL-4. The equilibrium dissociation constant of this binding interaction was found to be 459 nM. D) BLI association and dissociation curves for 500-4000 nM of Affibody 2 binding to 1 nM of biotinylated IL-4. The equilibrium dissociation constant of this binding interaction was found to be 141 nM.

**Table 2.**
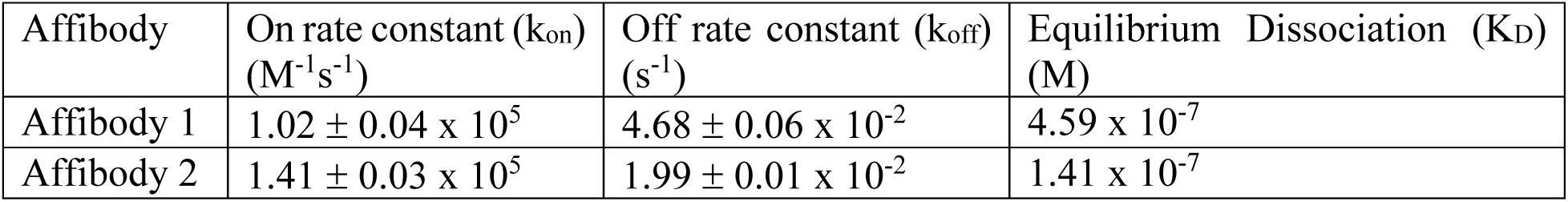
Binding kinetics of Affibody 1 and 2 interactions with IL-4.

| Affibody | On rate constant ($k_{on}$)<br>($M^{-1}s^{-1}$) | Off rate constant ($k_{off}$)<br>($s^{-1}$) | Equilibrium Dissociation ($K_D$)<br>(M) |
| --- | --- | --- | --- |
| Affibody 1 | $1.02 \pm 0.04 \times 10^5$ | $4.68 \pm 0.06 \times 10^{-2}$ | $4.59 \times 10^{-7}$ |
| Affibody 2 | $1.41 \pm 0.03 \times 10^5$ | $1.99 \pm 0.01 \times 10^{-2}$ | $1.41 \times 10^{-7}$ |

### Computational predictions of IL-4-specific affibody interactions with IL-4

To further explore the binding interactions between the IL-4-specific affibodies and IL-4, the molecular folding software AlphaFold3 and computational protein design software Rosetta were used to generate energy-minimized predictions of the bound structures of each affibody with IL-4 (**Figure 3A**). IL-4 presents as a 4-helix bundle, with distinct secondary structures identified as the A, B, C, and D helixes, which contribute to defined receptor binding motifs.^44,54–56^ AlphFold3 predicted that Affibody 1 would interact with IL-4 at the A and C helices,^57^ overlapping with the known binding interface for IL-4Rα, while Affibody 2 was predicted to interact with the BC loop, demonstrating no overlap with the IL-4Rα binding site (**Figure 3B**).^55^ Interface analysis of the IL-4 specific-affibodies demonstrated an enrichment of polar contacts at the Affibody 1 binding interface with IL-4 compared to the Affibody 2 binding interface. Neither of the affibody interactions with IL-4 were expected to overlap with the known binding interfaces of IL-4 with the γ-complex of another IL-4Rα or with IL-13Rα, which both occur at the AD face of IL-4.^55^

**Figure 3.**
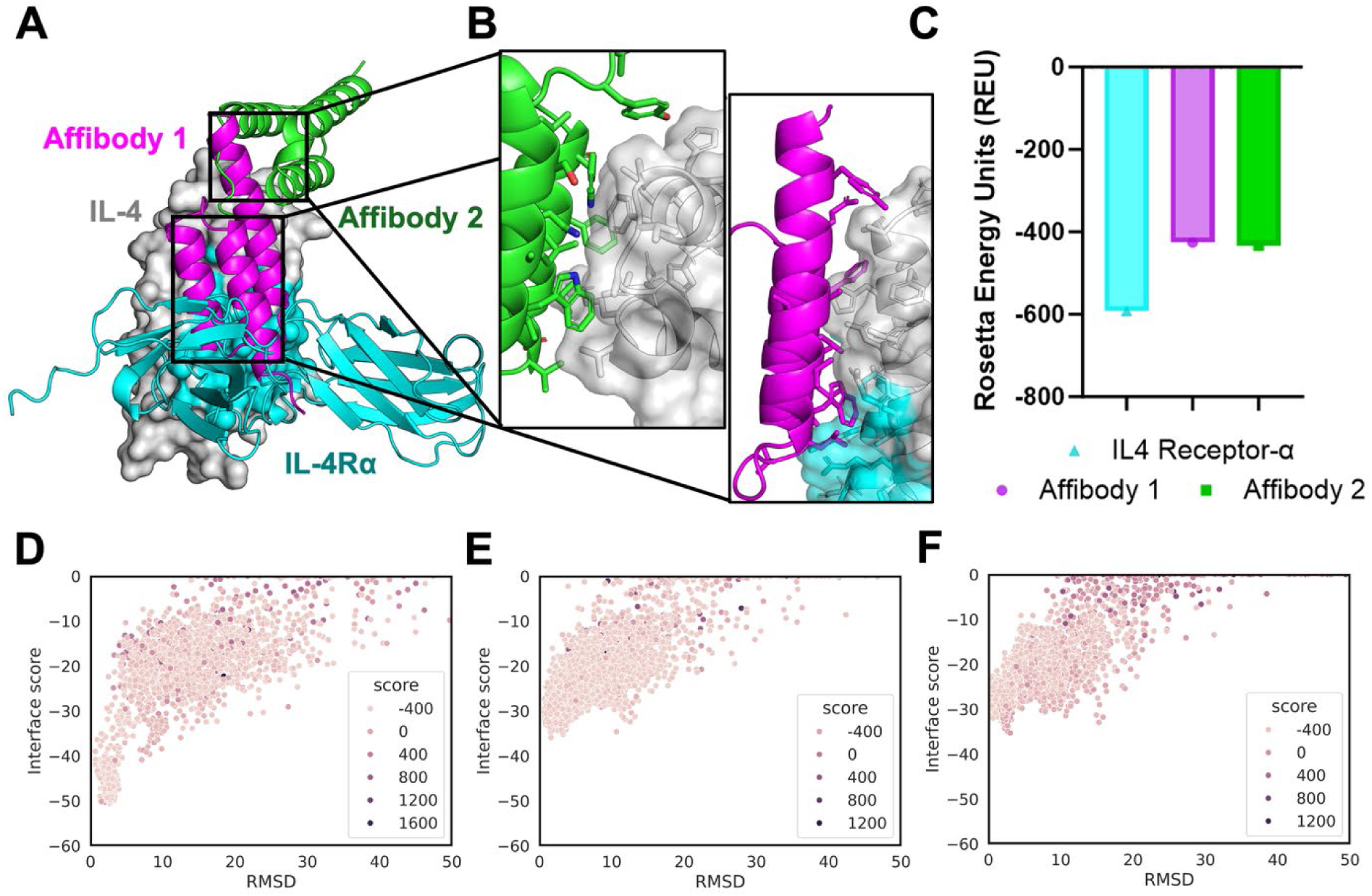
Computational predictions of IL-4-specific affibodies binding to IL-4. A) Molecular modeling of IL-4Rα (blue), Affibody 1 (pink), and Affibody 2 (green) docked onto IL-4 (white), as predicted by AlphaFold3 docking followed by Rosetta FastRelax energy minimization. B) Zoomed in interface analysis of Affibody 1 and Affibody 2 binding interactions with IL-4. C) Rosetta FastRelax bound state energies for IL-4 binding to IL4Rα, Affibody 1, and Affibody 2. RosettaDock interface scores of IL-4 binding to D) IL-4Rα, E) Affibody 1, and F) Affibody 2 as a function of RMSD from the initial modeled interfaces (n=5000).

Rosetta FastRelax was then used to quantify the binding energy for energy-minimized interactions between each affibody and IL-4 as well as the native IL-4-IL4Rα binding interaction.^44,46^ The binding interactions of Affibody 1 and Affibody 2 to IL-4 yielded comparable interface scores at the A and C helices and BC loop, respectively. Higher Rosetta Energy Unit (REU) interface scores were observed for both IL-4-specific affibodies binding to IL-4 compared to IL-4Rα receptor binding to IL-4, suggesting weaker interface interactions for the affibodies at the modeled sites than IL-4-IL4Rα binding (**Figure 3C**). RosettaDock was applied to further explore the interface specificity and binding energies of IL-4 interactions with IL-4-specific affibodies. Binding funnels were generated for IL-4Rα (**Figure 3D**), Affibody 1 (**Figure 3E**), and Affibody 2 (**Figure 3F**) binding to IL-4, suggesting equally stable binding interactions within 3 Å of the epitopes that were initially modeled and in agreement with the REUs predicted by Rosetta FastRelax. These computational predictions confirmed the BLI results, which suggest that the IL-4-specific affibodies engage in weaker interactions with IL-4 than the interaction between IL-4 and the IL-4Rα receptor.

Molecular dynamics simulations were performed to investigate the dynamics of binding between the IL-4-specific affibodies and IL-4. Starting from Rosetta energy-minimized IL-4-affibody bound structures, we performed quadruplicate 100,000 picosecond simulations of IL-4 binding to Affibody 1 and Affibody 2. We then modeled the α-carbon RMSD, affibody RMSF, ΔSASA_Total_, ΔSASA_Polar_, and ΔSASA_Nonpolar_ to explore differences in the predicted dynamics and solvent accessibilities of the interactions between IL-4 and the IL-4-specific affibodies.

Over 100,000 ps, Affibody 2 demonstrated an average α-carbon RMSD shift ranging between 3 and 7.5 Å with an average of approximately 4 Å. Conversely, Affibody 1 demonstrated a lower, more consistent RMSD shift ranging from 3 to 4 Å with an average of approximately 3.5 Å (**Figure 4A**). The comparatively small range of the α-RMSD of the interaction between IL-4 and Affibody 1 across quadruplicate trajectories suggest that interface elements of the Affibody 1 interaction with IL-4 limited the dynamics of the system and that a stable bound interface was achieved across each trajectory over the entire simulation. This result was further confirmed by comparison of the per residue RMSF for Affibody 1 and Affibody 2 contacting IL-4. Affibody 1 demonstrated a smaller 2.5 Å RMSF for interface contacting residues 10-30 on IL-4, and Affibody 2 demonstrated a larger 5-10 Å RMSF for the same residues on IL-4 (**Figure 4B, 4C**), suggesting that Affibody 1 underwent a smaller change in position at its binding interface with IL-4 over the simulation trajectory. No difference in ΔSASA_Total_ or ΔSASA_Nonpolar_ were observed over the trajectories of IL-4 binding to Affibody 1 or Affibody 2 (**Figure 4D, 4F**). However, Affibody 1 displayed a greater total ΔSASA_Polar_ than Affibody 2 (**Figure 4E**). These results may suggest that an enrichment of polar contacts at the binding interface between IL-4 and Affibody 1 is responsible for stabilizing the binding interaction over the simulated trajectories.

**Figure 4.**
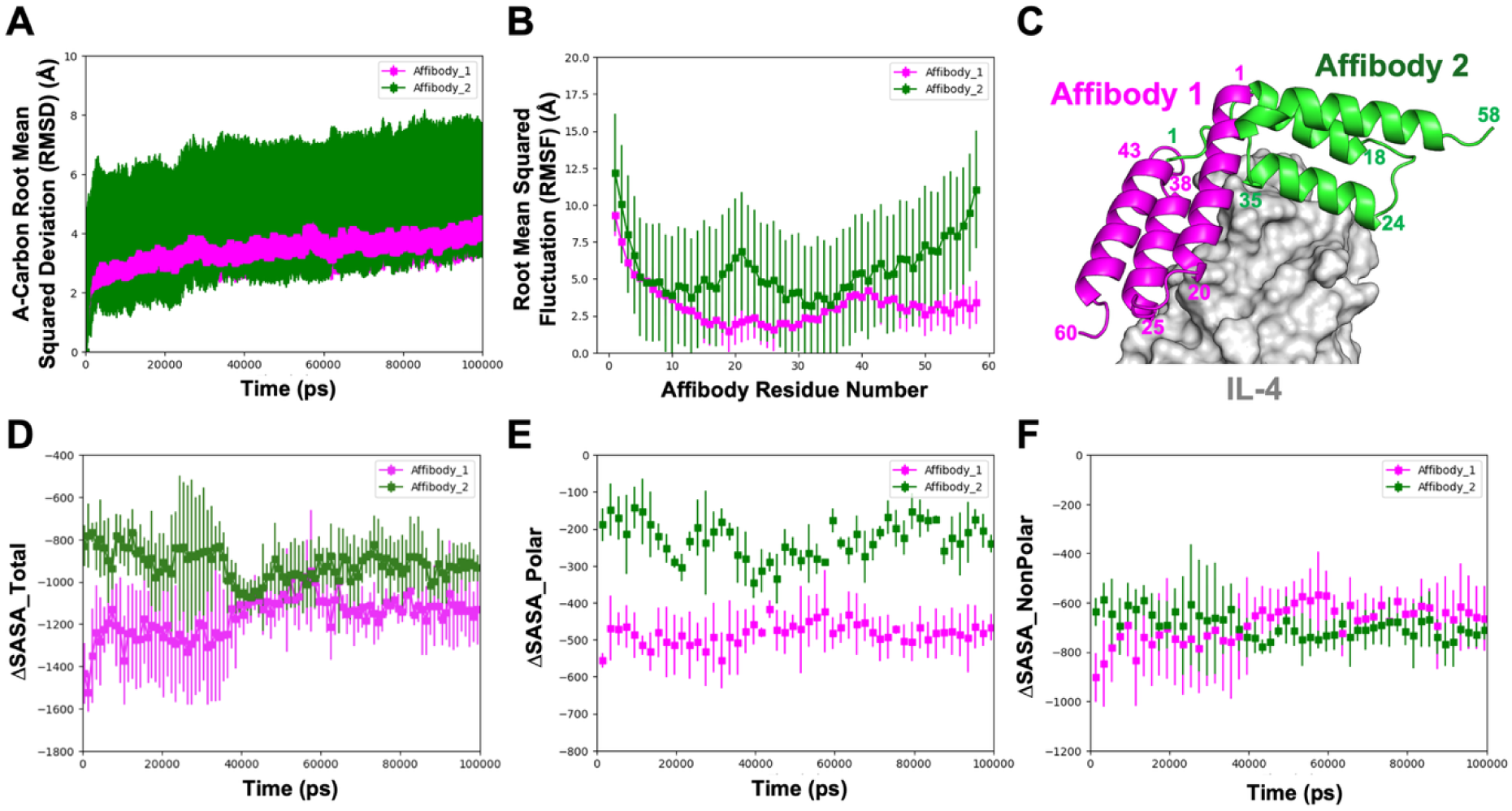
Molecular dynamics simulations of IL-4-specific affibody interactions with IL-4. A) All α-carbon RMSD of IL-4 binding to Affibody 1 or Affibody 2 over 100,000 ps (n=4). B) Per residue RMSF of Affibody 1 or Affibody 2 interacting with IL-4 over the 100,000 ps trajectory (n=4). C) Molecular model of Affibody 1 and Affibody 2 binding to IL-4 with residue numbers labeled. Change in the D) ΔSASA Total, E) ΔSASA Polar, and F) ΔSASA Nonpolar for Affibody 1 and Affibody 2 binding to IL-4 over 100,000 ps (n=4).

Taken together, the molecular dynamics simulations predicted that Affibody 1 underwent highly repeatable, less dynamic interactions with IL-4 than Affibody 2 across quadruplicate trajectories. Furthermore, the RMSF, ΔSASA_Total_, ΔSASA_Polar_, and ΔSASA_Nonpolar_ measurements suggest that this decrease in binding dynamics of the Affibody 1 interaction with IL-4 may be explained by a greater number of polar contacts at the its binding interface compared to IL-4-Affibody 2 binding, resulting in the formation of a more tightly packed, stable bound state.

### IL-4-specific affibodies modulate macrophage polarization

One of the critical functions of IL-4 is its role in the M2 polarization of macrophages. Thus, we chose to evaluate the effects of IL-4-specific affibodies on macrophages cultured with and without IL-4 to assess their impact on IL-4 bioactivity. Murine iBMDMs were cultured with IL-4, IL-4-specific affibodies, and IL-4 complexed with 5000 times molar excess of IL-4-specific affibodies for 3 days. Phase contrast images of iBMDMs treated with IL-4 or affibody-complexed IL-4 demonstrate that cells remained viable through the entire assessment period (**Figure 5A**). Gene expression of the M2-associated marker *Ym1* was evaluated, since its expression is a hallmark of IL-4 induced M2 macrophage polarization. As expected, exposure to IL-4 led to an approximately 10-fold increase in *Ym1* gene expression (**Figure 5B**). When IL-4 was complexed with either Affibody 1 or Affibody 2, *Ym1* gene expression trended slightly lower than IL-4 alone, although this difference was not statistically significant.

**Figure 5.**
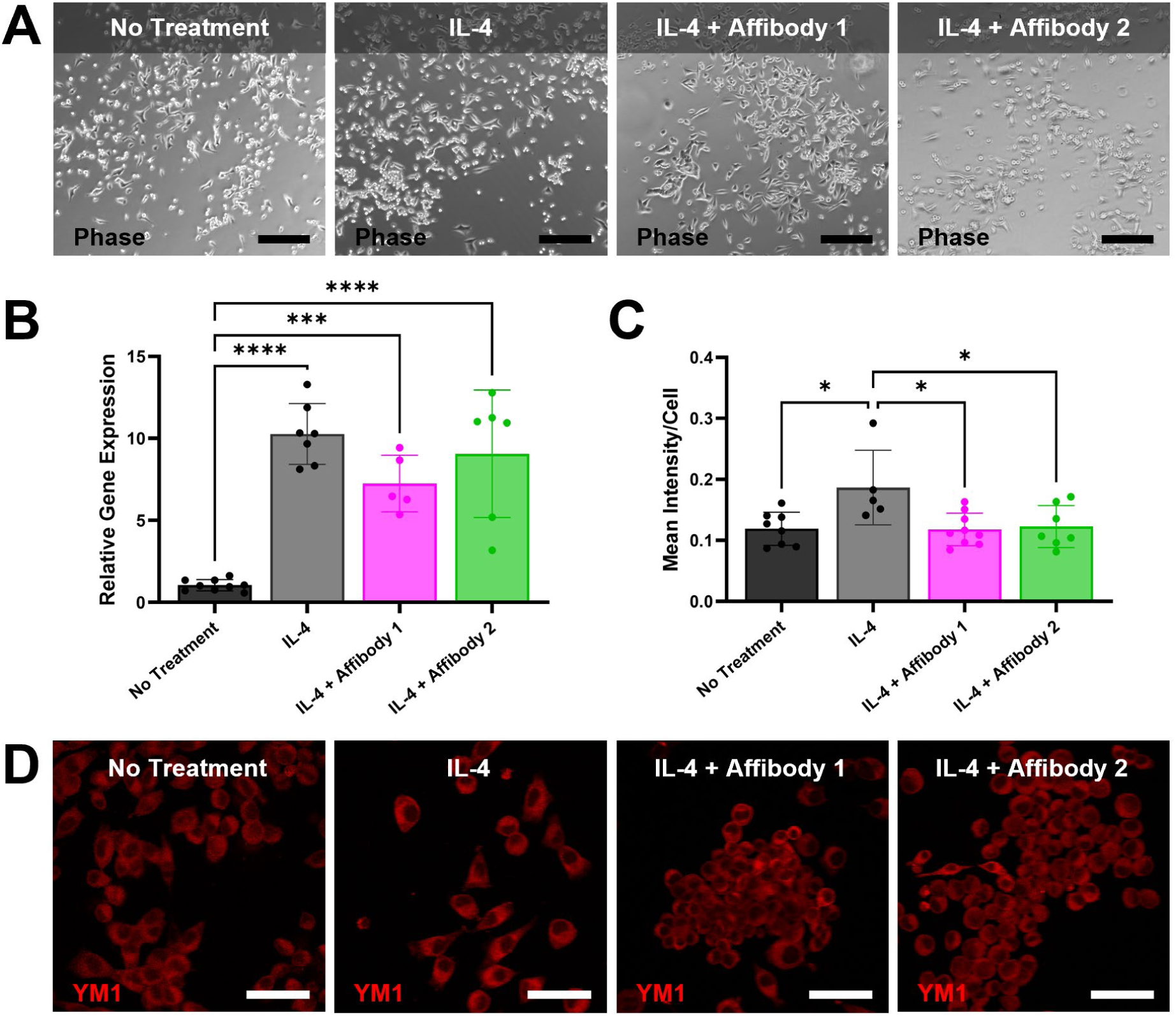
Effect of complexing IL-4 with IL-4-specific affibodies on Ym1 expression in bone marrow-derived macrophages. iBMDMs were cultured with no treatment, treatment with IL-4 (50 ng/mL), or treatment with 50 ng/mL of IL-4 complexed with Affibody 1 or Affibody 2 at a ratio of 1:5000. A) Representative phase contrast images of macrophages. Scale bar = 250 µm. B) Gene expression analysis of the *Ym1* gene. C) Quantification of Ym1 protein expression from confocal microscopy images, using mean Ym1 fluorescent intensity per cell. Statistical significance between groups was determined using one-way ANOVA with Tukey’s post-hoc test. * p < 0.05, *** p < 0.001, **** p < 0001, as indicated. n = 5-9. D) Representative confocal microscopy images of macrophages stained with anti-Ym1 antibody (red fluorescence). Scale bar = 20 µm.

At the protein level, immunofluorescence analysis revealed a similar, significant increase in Ym1 expression in response to IL-4 treatment (**Figure 5C, D**). Notably, this IL-4-induced increase was significantly reduced when IL-4 was complexed with Affibody 1 or Affibody 2. In contrast, treatment with Affibody 1 or Affibody 2 alone did not alter Ym1 levels (**Figure S1**), suggesting that IL-4 polarizes macrophages toward an M2 phenotype, and Affibody 1 or Affibody 2 inhibit the ability of IL-4 to induce M2 polarization and Ym1 expression.

### Controlled release of IL-4 from affibody-conjugated hydrogels

100 µL 5% (w/v) affibody-conjugated PEG-mal hydrogels were loaded with 100 ng of IL-4 overnight and allowed to release IL-4 over 7 days into a solution of 10% FBS in PBS. Affibody 1 and Affibody 2 were conjugated to the hydrogels at 100, 500, and 5000 molar excesses to IL-4 to control protein release. All hydrogels encapsulated more than 90% of the loaded IL-4 (**Figure 6A, 6D**); however, IL-4 loading was significantly lower in hydrogels without affibodies compared to hydrogels with all molar excesses of Affibody 1 as well as the highest molar excess of Affibody 2. As expected, PEG-mal hydrogels without affibodies released almost all of the loaded IL-4 over 7 days (**Figure 6B, 6E**). In contrast, affibody conjugation to the hydrogels significantly reduced IL-4 release over the same time period. There was a dose-dependent effect of affibody conjugation on IL-4 release. Hydrogels containing 5000 times molar excess of either affibody released less IL-4 than other affibody-conjugated hydrogels at most time points, and additional significant differences in IL-4 release were observed between hydrogels containing 100 and 500 times molar excess of affibodies at several time points. After 7 days, 90.62 ± 3.85% of the loaded IL-4 was recovered from the hydrogels without affibodies, while only 10.65 ± 0.83% and 8.64 ± 0.3% of the loaded IL-4 were recovered from the hydrogels containing the highest molar excess of Affibody 1 and Affibody 2, respectively. By plotting the fractional protein release against the square root of time, IL-4 release was determined to follow Fickian diffusion.^58^ Based on the slopes of the linear portions of the Fickian diffusion plots (**Figure S2**), hydrogels without affibodies released IL-4 significantly faster than all hydrogels containing Affibody 1 or Affibody 2 (**Figure 6C, 6F**). There was also a dose-dependent effect of affibody amount on IL-4 release rate, as 5000 molar excess of either affibody significantly slowed the release rate of IL-4 compared to 100 and 500 molar excess of the affibodies. No differences were observed in IL-4 loading or release between hydrogels conjugated with the same amounts of Affibody 1 or Affibody 2. Overall, the incorporation of IL-4-specific affibodies increased IL-4 loading and decreased the amount and rate of IL-4 release in a dose-dependent manner, demonstrating the ability of IL-4-specific affibodies to finely tune the delivery and presentation of IL-4 from PEG-mal hydrogels.

**Figure 6.**
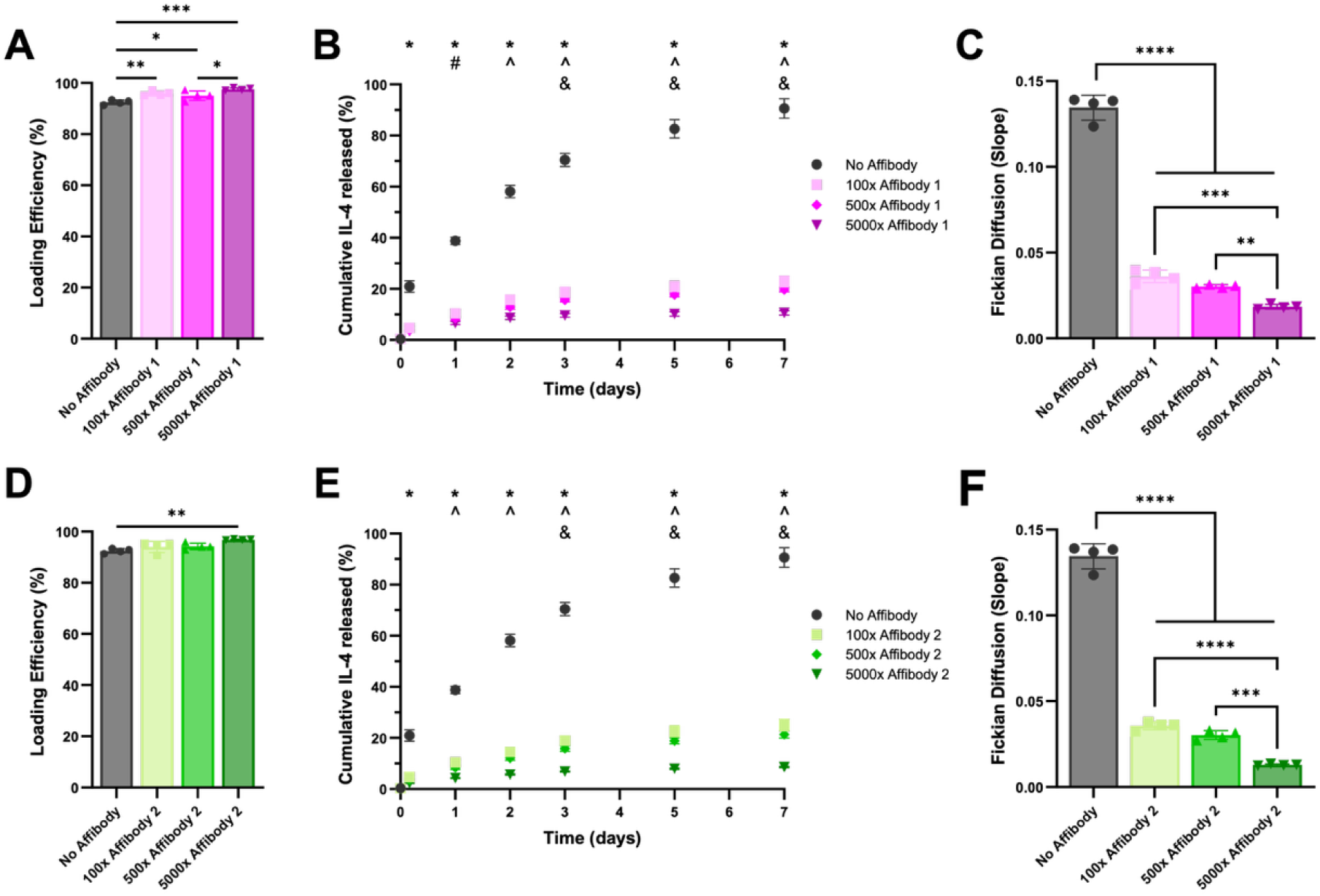
Release of IL-4 from affibody-conjugated PEG-mal hydrogels. PEG-mal hydrogels with and without IL-4 specific affibodies were incubated with 100 ng of IL-4 and allowed to release protein into 10% (v/v) FBS in PBS over 7 days. Percentage of initial IL-4 loaded in hydrogels conjugated with A) Affibody 1 and C) Affibody 2 at different molar excesses to IL-4 (100x, 500x, 5000x) compared to hydrogels without affibodies. Statistical significance was determined using 1-way ANOVA followed by Tukey’s post-hoc test. * p < 0.05, ** p < 0.01, *** p < 0.001, as indicated. Cumulative release of loaded IL-4 from hydrogels conjugated with B) Affibody 1 and E) Affibody 2 at different molar excesses to IL-4 (100x, 500x, 5000x) compared to hydrogels without affibodies. Statistical significance was determined using 2-way ANOVA followed by Tukey’s post-hoc test. * p < 0.05 for PEG Only vs. all other groups, ^ p < 0.05 for 5000x Affibody vs. all other molar excesses of affibodies, # p < 0.05 for 5000x Affibody vs. 100x Affibody, and & p < 0.05 for 100x Affibody vs. 500x Affibody. Fickian diffusion rates of IL-4 from PEG-mal hydrogels conjugated with C) Affibody 1 and F) Affibody 2 at different molar excesses to IL-4 (100x, 500x, 5000x) compared to hydrogels without affibody, indicated by the slopes of cumulative release plotted against the square root of time (Figure S2). Statistical significance was determined using 1-way ANOVA followed by Tukey’s post-hoc test. ** p < 0.01, *** p < 0.001, **** p < 0.0001, as indicated. n = 4.

## Discussion

IL-4 is a potent immunoregulatory cytokine with the ability to modulate the activities of several immune cell populations, giving it promising therapeutic potential. However, its development for clinical applications has been hindered by the challenge of achieving sustained delivery that can be tuned to provide the appropriate temporal dynamics to resolve inflammation without causing immune dysregulation. We identified two IL-4-specific affibodies that can be used to modulate IL-4 activity and integrated into a biomaterial platform to provide sustained IL-4 delivery. By changing the ratio of affibodies to IL-4, we demonstrated that we could tune the amount and rate of IL-4 released from PEG-mal hydrogels.

While other strategies have been developed for sustained IL-4 delivery, few biomaterial delivery vehicles have demonstrated the degree of tunability afforded by protein-specific affinity interactions. Affinity-controlled release of IL-4 was recently explored in another study, in which biotinylated IL-4 was released from hydrogels modified with streptavidin, avidin, or CaptAvidin.^19^ By leveraging the strong affinity interactions between biotin and avidin, IL-4 release could be tuned by using different avidin variants and accelerated by adding excess free biotin. Interestingly, the avidin variants themselves also affected macrophage phenotype. Comparatively, the interactions between IL-4-specific affibodies and IL-4 are significantly weaker than biotin-avidin interactions (K_D_ = 10^−7^ M vs. K_D_ = 10^−15^ M), yet do not require protein modification and are stable in the presence of serum proteins, making them advantageous for *in vivo* applications that require a substantial amount of IL-4 released over a sustained period of time. We also did not find IL-4-specific affibodies to affect Ym1 expression and M2 polarization of murine macrophages in the absence of IL-4. These results corroborate results from other studies, in which affibodies specific for bone morphogenetic protein-2 (BMP-2), vascular endothelial growth factor (VEGF), and platelet-derived growth factor (PDGF) were not found to impact the expected functional outcomes of these growth factors on their own, suggesting that affibodies may not elicit substantial off-target effects.^26,50^

Because affibodies discovered using yeast surface display libraries have been predicted to bind to receptor binding epitopes on the target protein,^26,50^ we were interested in further exploring the binding interfaces between IL-4 and the IL-4-specific affibodies *in silico* to predict how affibody binding to IL-4 would impact the ability of IL-4 to interact with its receptors. Affibody 1 was predicted to bind to IL-4 at the IL-4Rα binding site, and Affibody 2 was predicted to bind in another location distal to the IL-4Rα, IL-4Rα γ, and IL-13Rα binding interfaces on IL-4. Thus, we originally expected that Affibody 1 binding to IL-4 would impact macrophage responses downstream of IL-4 signaling more strongly, and Affibody 2 binding to IL-4 would have minimal effects on IL-4 activity. However, we found that both IL-4-specific affibodies significantly decreased IL-4-activated Ym1 protein expression, with only minimal changes in *Ym1* gene expression. These results may be explained by the fact that IL-4 signaling in macrophages requires secondary binding of IL-4 to an additional IL-4Rα or IL-13Rα to form the IL-4 receptor complex. Molecular dynamics simulations indicated that the binding of IL-4 to multiple receptors may be sterically hindered by Affibody 1 binding. Thus, IL-4-specific affibodies may have additional utility as IL-4 inhibitors. Current methods to block IL-4 signaling, such as antibodies (e.g., pascolizumab^59^) and soluble receptors (e.g., altrakincept^60^), have limited efficacy and require frequent dosing.^61^ IL-4-specific affibodies may overcome some of these limitations, especially when combined with inhibitors for the redundant IL-13 pathway.

Lastly, our use of yeast surface display libraries to discover affibodies for other target proteins have yielded multiple protein-specific affibodies with a range of affinities for tuning protein release.^26,50,62^ The use of this method with IL-4 required more sorting steps and only identified two unique IL-4-specific affibodies with similar affinities for IL-4. Thus, to achieve tunability of IL-4 release, we opted to change the molar excess of affibodies to IL-4, changing the avidity of the interactions instead of the affinity. This approach successfully provided different rates and amounts of IL-4 release that were all significantly lower than the burst release of IL-4 exhibited from PEG-mal hydrogels without affibodies. In the future, the release kinetics of IL-4 from these hydrogels could be further tuned by generating additional IL-4-specific affibodies with novel affinities by mutating the existing IL-4-specific affibodies using site-directed mutagenesis or computational protein design.^26^

## Conclusion

In conclusion, this work is the first demonstration of affinity-based IL-4 delivery using IL-4-specific affibodies, marking a significant advance in the development of controlled delivery vehicles for IL-4. Although IL-4 affibodies were predicted to bind to different locations on IL-4, they elicited the same effects on M2 polarization of murine macrophages, resulting in inhibition of IL-4-induced Ym1 protein expression. IL-4 delivery from PEG-mal hydrogels containing either IL-4-specific affibody could be tuned by changing the number of affibodies conjugated to the hydrogel relative to the amount of IL-4 initially loaded. Future work will involve further investigating the effects of IL-4 released from affibody-conjugated hydrogels and IL-4 complexed with affibodies on additional aspects of macrophage function and other immune cell populations involved in orchestrating the immune response.

## Acknowledgements

We are grateful for funding from the National Institutes of Health (R35 Maximizing Investigators’ Research Award (MIRA) R35-GM147507), Department of Defense (Discovery Award, Peer-Reviewed Medical Research Program W81XWH2210700 to M.H.H. and HT9425-24-1-0998 through the Toxic Exposures Research Program to S.A.R.), and Lary Simpson Professorship to M.H.H. J.D. was supported by a doctoral-level post-graduate scholarship (PGS-D) from the Natural Sciences and Engineering Research Council (NSERC) of Canada. J.E.S. is currently supported by an NIH Ruth L. Kirschstein Predoctoral Individual National Research Service Award (F31HL176164) and was previously supported by an NSF Research Traineeship in Molecular Probes and Sensors for Complex Environments (2022168). The yeast surface display library was a gift from Dr. Benjamin Hackel’s laboratory at the University of Minnesota (Minneapolis, MN), and the murine immortalized bone marrow-derived macrophages were a gift from Dr. Philip West’s laboratory at Jackson Labs (Bar Harbor, ME). The scripts used to set up and run our simulations were provided by Dr. Michael J Harms and are freely available at https://github.com/harmslab/setup_md. We thank members of the Hettiaratchi Lab for their thoughtful review of this manuscript. We also thank Dr. Danielle Benoit for the use of her lab’s peptide synthesizer and HPLC system.

