## Supplementary Document for "Discovery and Characterization of Interleukin-4-Specific Affibodies for Affinity-Controlled Protein Release and Macrophage Polarization"

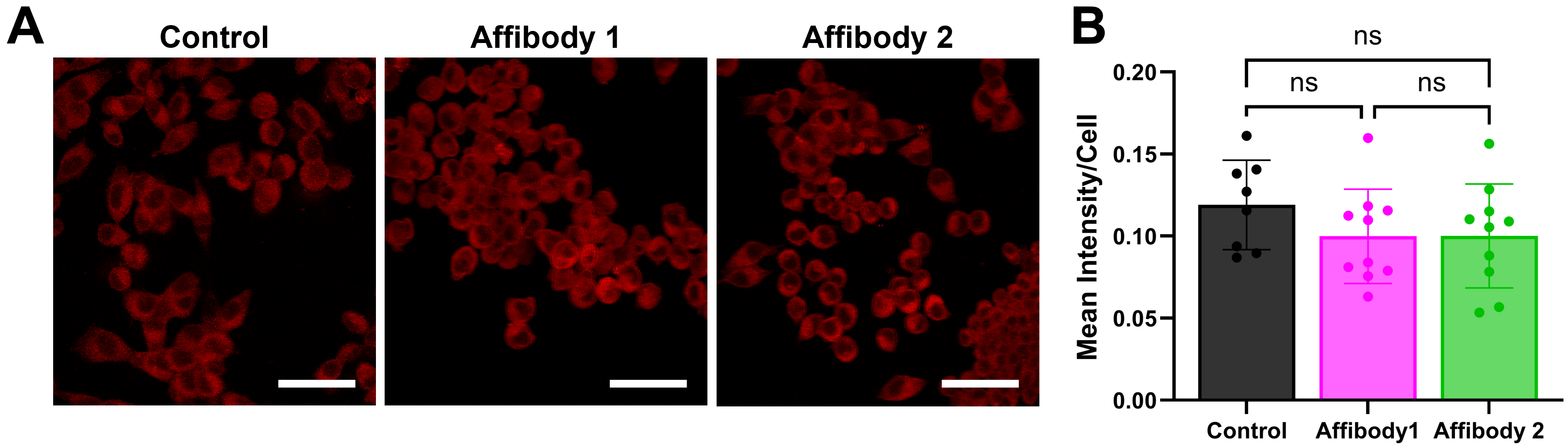

**Figure S1. Effect of IL-4-specific affibodies on Ym1 expression in bone marrow-derived macrophages.** iBMDMs were cultured with no treatment or treatment with 250 μg/mL of Affibody 1 or Affibody 2. A) Representative confocal microscopy images of macrophages stained with anti-Ym1 antibody (red fluorescence). Scale bar = 20 µm. B) Quantification of Ym1 protein expression from confocal microscopy images, using mean Ym1 fluorescent intensity per cell. Statistical significance between groups was determined using one-way ANOVA with Tukey’s post-hoc test. n = 8-10. ns = not significant.

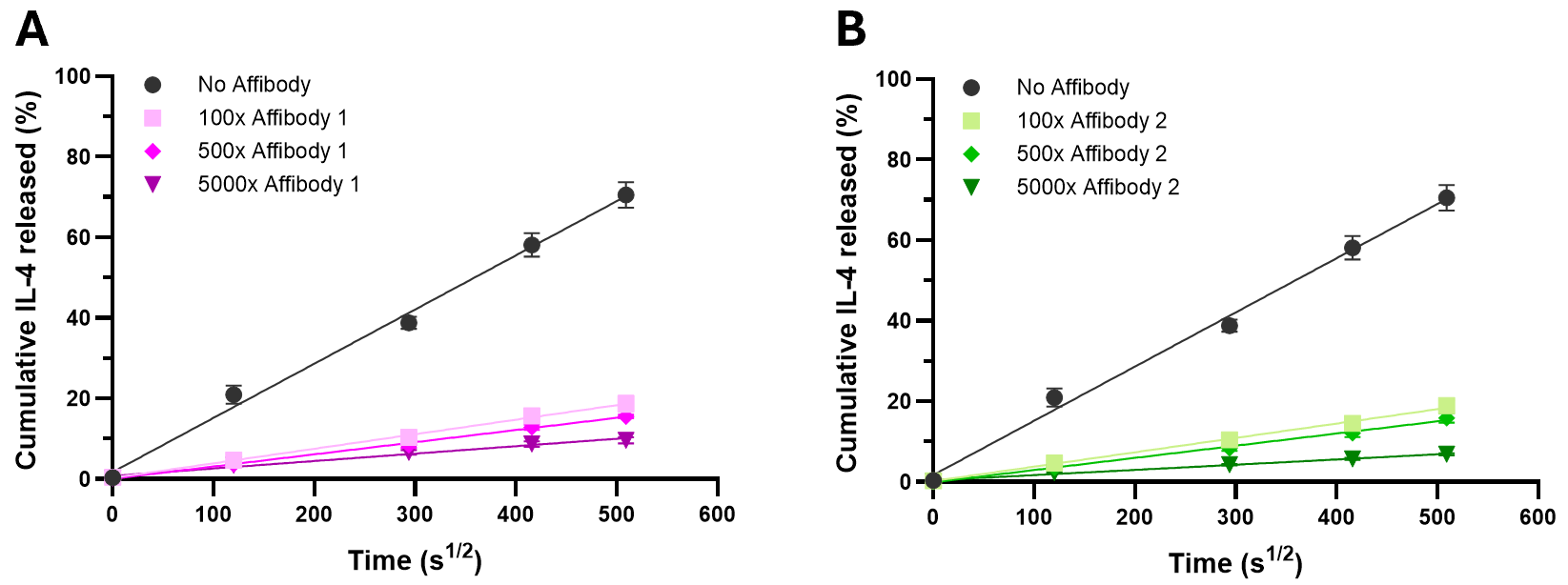

**Figure S2. Cumulative IL-4 release plotted as a function of the square root of time, representative of Fickian diffusion.** PEG-mal hydrogels with and without IL-4 specific affibodies were incubated with 100 ng of IL-4 and allowed to release protein into 10% (v/v) FBS in PBS over 7 days. Cumulative release of loaded IL-4 from hydrogels conjugated with A) Affibody 1 and B) Affibody 2 at different molar excesses to IL-4 (100x, 500x, 5000x) compared to hydrogels without affibodies and plotted again the square root of time. The slope of the linear portion of the release profile is representative of effectivity diffusivity when the release profile follows Fickian diffusion. n = 4.
